# Structural brain imaging predicts individual-level task activation maps using deep learning

**DOI:** 10.1101/2020.10.05.306951

**Authors:** David G. Ellis, Michele R. Aizenberg

## Abstract

Accurate individual functional mapping of task activations is a potential tool for biomarker discovery and is critically important for clinical care. While structural imaging does not directly map task activation, we hypothesized that structural imaging contains information that can accurately predict variations in task activation between individuals. To this end, we trained a convolutional neural network to use structural imaging (T1-weighted, T2-weighted, and diffusion tensor imaging) to predict 47 different functional MRI task activation volumes across seven task domains. The U-Net model was trained on 654 subjects and then subsequently tested on 122 unrelated subjects. The predicted activation maps correlated more strongly with their actual maps than with the maps of the other test subjects. An ablation study revealed that a model using the shape of the cortex alone or the shape of the subcortical matter alone was sufficient to predict individual-level differences in task activation maps, but a model using the shape of the whole brain resulted in markedly decreased performance. The ablation study also showed that the additional information provided by the T2-weighted and diffusion tensor imaging strengthened the predictions as compared to using the T1-weighted imaging alone. These results indicate that structural imaging contains information that is predictive of inter-subject variability in task activation mapping and cortical folding patterns as well as microstructural features may be a key component to linking brain structure to brain function.

## Introduction

Functional MRI (fMRI) maps the locations and intensities of brain activations by measuring the blood-oxygen-level-dependent (BOLD) signal arising from task performance in the scanner. In a research setting, the correlation between the performed task and subsequent BOLD response is estimated for the cortex and averaged over a group of subjects. Mapping brain task function gives researchers insight into regions of interest that activate during task performance for the group being studied. Non-invasive mapping of task activations in the brain using fMRI has also been appealing for research and clinical use in individual subjects. Accurate task activation mapping of individuals is critically important in patient care, such as brain tumor cases requiring neurosurgical resection. Research has shown, however, that individual task fMRI (tfMRI) activation maps have both limited reliability and accuracy (Elliott et al., 2020; Ellis et al., 2020; Weng et al., 2018). Therefore, investigation into alternatives for individual task mapping is warranted.

One potential alternative to relying solely on tfMRI for individual subject mapping is to deduce localization and intensity of task activations from structural imaging, such as anatomical and diffusion imaging. Unfortunately, there is little evidence demonstrating that structural imaging features can predict variations in task activations from multiple domains in individual subjects. Tavor et al. demonstrated that while a linear model using both structural and resting-state fMRI features could predict individual subject variations in task activation, an identical model using only structural features could not. The more complex method of extracting white matter tractography-based connectivity features from the diffusion signal has shown promise for predicting individual differences in responses to visual stimuli (Osher et al., 2015; Saygin et al., 2012; Saygin et al., 2016), but the same has not been reported for a broader range of task domains.

We hypothesize that a trained convolutional neural network (CNN) will predict task activation maps from diffusion and anatomical imaging that are sensitive to inter-subject differences over a wide variety of task domains. This finding would demonstrate that structural imaging features do contain variances predictive of functional task differences in individuals.

## Methods

### Data Acquisition

Imaging data was obtained from the Human Connectome Project (HCP) young adults S1200 public data release (https://www.humanconnectome.org/study/hcp-young-adult) provided by the WU-Minn HCP consortium. All subjects were 22 to 35 years old and were scanned by the HCP using Washington University’s 3T Siemens Connectome Scanner (D C Van Essen et al., 2012). Preprocessed and aligned anatomical and structural data was provided by the HCP in the subject T1-weighted (T1w) imaging space, including: T1w and T2-weighted (T2w) with 0.7mm isotropic voxel size, diffusion images with 1.25mm isotropic voxel size acquired at 3 b-shells (1000, 2000, and 3000 s/mm^2^) with 90 directions per shell acquired twice with opposite phase encoding directions (Sotiropoulos et al., 2013), and MSMSulc and MSMAll (Glasser et al., 2016) registered template surfaces. The HCP also provided two runs of unprocessed task fMRI volumes for seven task domains (Barch et al., 2013): hand, foot, and tongue movements (MOTOR), auditory language processing (LANGUAGE), n-back working memory (WM), shape and texture matching (RELATIONAL), emotion-processing (EMOTION), social interactions (SOCIAL), and incentive processing (GAMBLING).

### Subject Selection

Of the 1206 HCP subjects, 654 were selected for the training group, 133 for the validation group, and 122 unrelated subjects for the test group. The remaining 297 subjects were excluded for having incomplete data, failing data processing, or being related to the test group. In order to prevent information from one of the groups biasing the results of another group, all groups were selected so that no subjects from one group were related to any of the subjects in another group. Additionally, 39 subjects that were scanned twice by the HCP with mean interval of 140 days were used to evaluate the test-retest reliability of the predicted and actual maps. These test-retest subjects were either a part of the test group or related to someone in the test group, and none of them were a part of the training or validation groups.

### MRI Processing

The fMRI volume preprocessing was performed according to the HCP processing pipelines including gradient distortion correction (Glasser et al., 2013), with the exception that the one-step resampling linearly transformed the volumes into the native T1w space rather than the MNI space. Individual-level fMRI z-score activation volumes were then computed across both runs for each task domain using the HCP pipelines in the subject’s native volume space with high-pass bandwidth filtering and minimal 2mm FWHM smoothing (Woolrich, Ripley, Brady, & Smith, 2001). All 47 unique task activation volumes across the seven task domains were used for model training and testing (*Table S1*). The preprocessed T1w, T2w, and diffusion data were used as distributed by the HCP. Diffusion tensor image (DTI) modeling was performed using dipy (Garyfallidis et al., 2014) on the preprocessed diffusion data resulting in mean diffusivity (MD) and 3-directional fractional anisotropy (FA) feature maps computed for each b value separately (1000, 2000, and 3000 s/mm^2^) in the subject’s native space similar to Ganepola (Ganepola et al., 2018).

### Model Architecture

For predicting task activation from structural imaging, a U-Net-style CNN was used (*Figure S1*) (Myronenko, 2018; Ronneberger, Fischer, & Brox, 2015). This model architecture combines learned convolutional layers at multiple resolutions. The initial input images consisted of the aligned and skull-stripped T1w, T2w, and DTI volumes cropped to remove background slices and resampled to a size of 144×160×144. Each layer consisted of two residual blocks (He, Zhang, Ren, & Sun, 2016) performing group normalization (Wu & He, 2018), rectified linear activation, a 3×3×3 convolution (Myronenko, 2018). The first encoding layer consisted of 32 channels and contained a 20% channel dropout between the two residual blocks. After each consecutive encoding layer the images were downsampled by a factor of two using a strided convolution. As is common in U-Net architectures, the following layer doubled the number of channels. The encoder consisted of five layers. Each layer of the decoder mirrored the encoder with two residual blocks per layer. The decoder layers took as inputs the upsampled output of the previous decoder layer concatenated with the output of the encoder layer of the same depth and resolution. A final 1×1×1 convolution linearly resampled the outputs from 32 channels of the last decoding layer to predict the output volumes for each of the 47 task activation maps. The predicted output volumes were then resampled back into the original T1w space.

### Model Training

We trained a single model to predict the fMRI task activation volumes across all seven task domains using T1w, T2w, and DTI data from the training set of subjects (*Figure 1A*). The mean squared error loss between the predicted and actual activation volumes was used to iteratively train the model. The loss function was weighted so that each of the seven domains had equal influence on the model training. In order to augment the training data, the scale of the input and output images was randomly altered, and random white noise was added to the input images. The learning rate was decayed after the validation loss had not improved for 20 epochs, and model training was stopped after the validation loss did not improve for 50 epochs.

**Figure 1.**
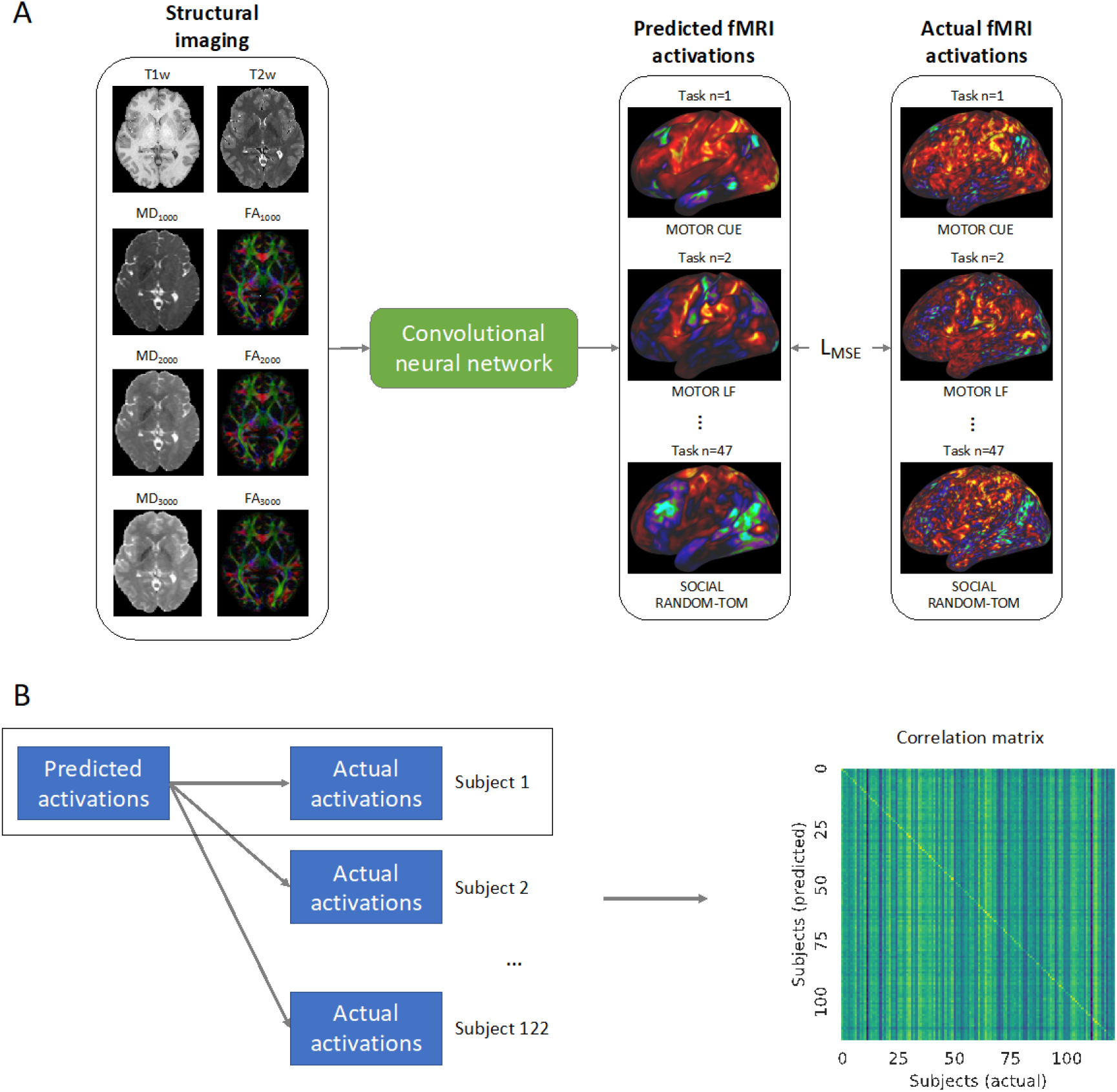
(**A**) Training of the convolutional neural network. The model takes as inputs the structural imaging for a given subject which includes the T1w and T2w weighted imaging as well as mean diffusivity (MD) and fractional anisotropy (FA) for b-values 1000, 2000, and 3000 s/mm^2^. The model is trained to predict the task activation volumes for 47 different task maps across 7 different task domains (visualized on the cortical surface in the figure). The mean squared error loss (LMSE) between the predicted and actual activations is used to iteratively train the model. (**B**) In order to validate the model’s ability to predict individual activation maps, the predicted maps of a given subject *m* are compared to the actual activation maps for that subject as well as the activation maps for all of the 122 subjects. From these comparisons, the correlation matrix shown on the right is constructed.

### Model Testing and Statistical Analysis

After the completion of model training, the model was used to predict the activation maps of 47 different tfMRI maps from seven different task domains. In order to validate the model’s ability to detect individual differences in task maps, we imitated the analysis done by Tavor et al. (Tavor et al., 2016) and compared the correlations between the predicted and actual maps (*Figure 1B*). To allow for the comparison between subjects, both the actual and predicted task activation volumes were sampled onto the cortical template surfaces in T1w space using the connectome workbench (https://humanconnectome.org/software/connectome-workbench) under a cortical ribbon constrained sampling method. The cortical template surfaces are distributed by the HCP with indices that have been aligned between subjects according to sulcal patterns (MSMSulc) using surface matching registration (Coalson, Van Essen, & Glasser, 2018). The predicted task activation maps of a given subject were then compared to the actual activation maps for that subject as well as to the activation maps for all of the other test subjects and a correlation matrix was constructed. A Kolmogorov-Smirnov test between the distribution of the diagonal elements of the correlation matrix and the extra-diagonal elements was performed with the threshold for significance set at *α* = .05.

In order to determine if the inter-subject variation predicted by the model could be accounted for via alignment with structural and functional features beyond that of the just the sulcal patterns, the predicted and actual activation volumes were sampled onto another set surface templates whose indices had been aligned by the HCP using the MSMAll surface matching technique (Coalson et al., 2018; Glasser et al., 2016). Correlations between the predicted and actual activations as sampled onto the MSMAll surface templates were computed and a correlation matrix was constructed. A Kolmogorov-Smirnov test was again conducted to test for a difference between the distributions of the diagonal and extra-diagonal elements.

The difference in the distributions was also tested for each of the 47 task activations with the threshold corrected for multiple comparisons: 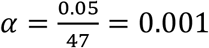. The lateralization indices (the difference in activation between the left hemisphere and right hemisphere averaged from the surface vertices within a 10mm radius of the peak location for the predicted map) as described by Tavor et al (Tavor et al., 2016) for the predicted maps were also compared to the lateralization indices for the actual maps and tested with a correlation analysis corrected for multiple comparisons (*α* = .001).

An ablation study was performed to assess the amount of information gained by the selected input features. In this study, in addition to the CNN model trained as described previously, six additional models were trained in an identical fashion but with different input feature maps: (1) using separately processed DTI features not that were not separated for each b-value (as was done in the main model) along with T1w and T2w imaging, (2) using T1w and T2w imaging alone, (3) T1w imaging alone, (4) a binary mask of the cortex, (5) a binary mask of the subcortical matter, and (6) a binary mask of the whole brain. In order to observe the effects of alignment techniques, the ablation study was evaluated using the MSMSulc and MSMAll template surfaces, as well as with the predicted and activation volumes aligned via non-linear warping into MNI space.

Additionally, a test-retest reliability analysis was performed on the actual maps and predictions by calculating the intraclass correlation (ICC) using the ICC(3,1) mixed-effects model (Shrout & Fleiss, 1979).

## Results

The CNN was able to use structural imaging to predict activation patterns of individual subjects that matched the group average as well as activations that deviated from the group average, (*Figure 2*). The predicted maps matched their actual maps better than the maps of other subjects, as seen by the diagonal dominance correlation matrix (*Figure 3A*). Row and column normalization to remove the mean and to account for the higher variability in the actual maps made diagonal of the correlation even more pronounced (*Figure 3B*). A Kolmogorov-Smirnov test between the distribution of the diagonal elements of the correlation matrix and the extra-diagonal elements (*Figure 3C*) resulted in a highly significant difference (*p* < .001) indicating that the predicted task maps are significantly more correlated to their actual maps than to the maps of other subjects. Likewise, the distributions of all of the individual task maps passed the test score corrected for multiple comparisons of (*p* < .001) except for the GAMBLING Punish-Reward task map (*p* = .23), which is known to have limited utility, and the MOTOR LF-AVG (left foot minus average, *p* = .009) task map (*Figures S2-S4*). For all task maps, the average correlation to its actual map was greater than the average correlation to the actual maps of the other subjects, showing that, on average, the predicted maps matched their actual maps more than the mean (*Figures S5*). Even with using the MSMAll registered template surfaces the Kolmogorov-Smirnov test between the distribution of the diagonal elements of the correlation matrix and the extra-diagonal elements resulted in a highly significant difference (*p* < .001) (*Figure S6*). Furthermore, all predicted maps still matched their actual maps better than the other actual maps, on average, for all domains and input types when using the MSMAll registered template surfaces (Figure 4).

**Figure 2.**
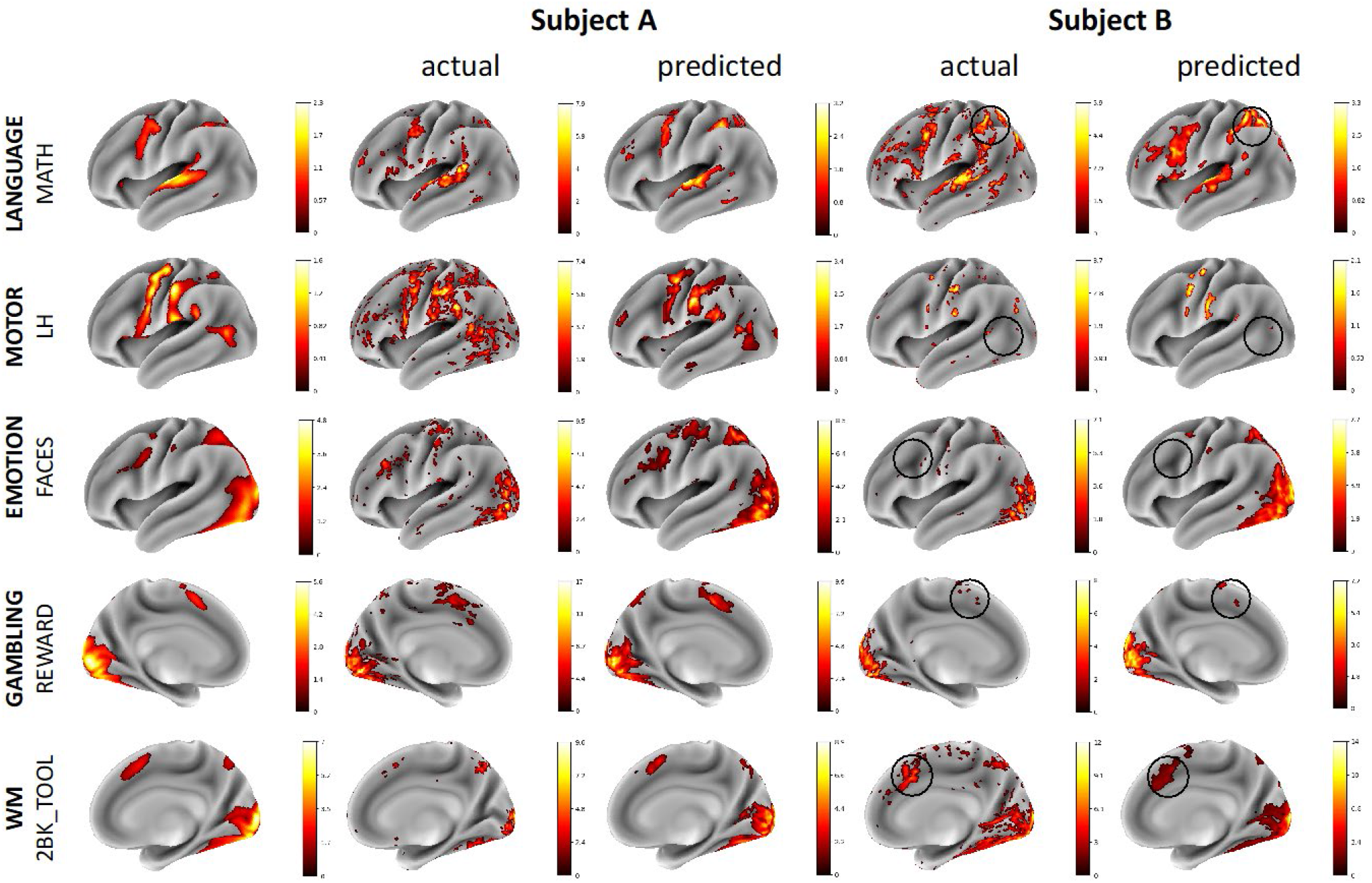
Example comparisons between group average, actual, and predicted maps for illustrative subjects with task maps across several domains. The actual and predicted Subject A maps demonstrate the model’s ability to predict accurate activation maps that resemble the group average while the Subject B maps demonstrate the model’s ability to predict deviations from the group average. Thresholds for the maps were determined using the medians of the positive and negative gamma distributions from a Gaussian and 2-gamma mixture model. The black circles highlight variations from the group average that the model was able to predict correctly. Subjects A and B are different subjects for each of the various task maps.

**Figure 3.**
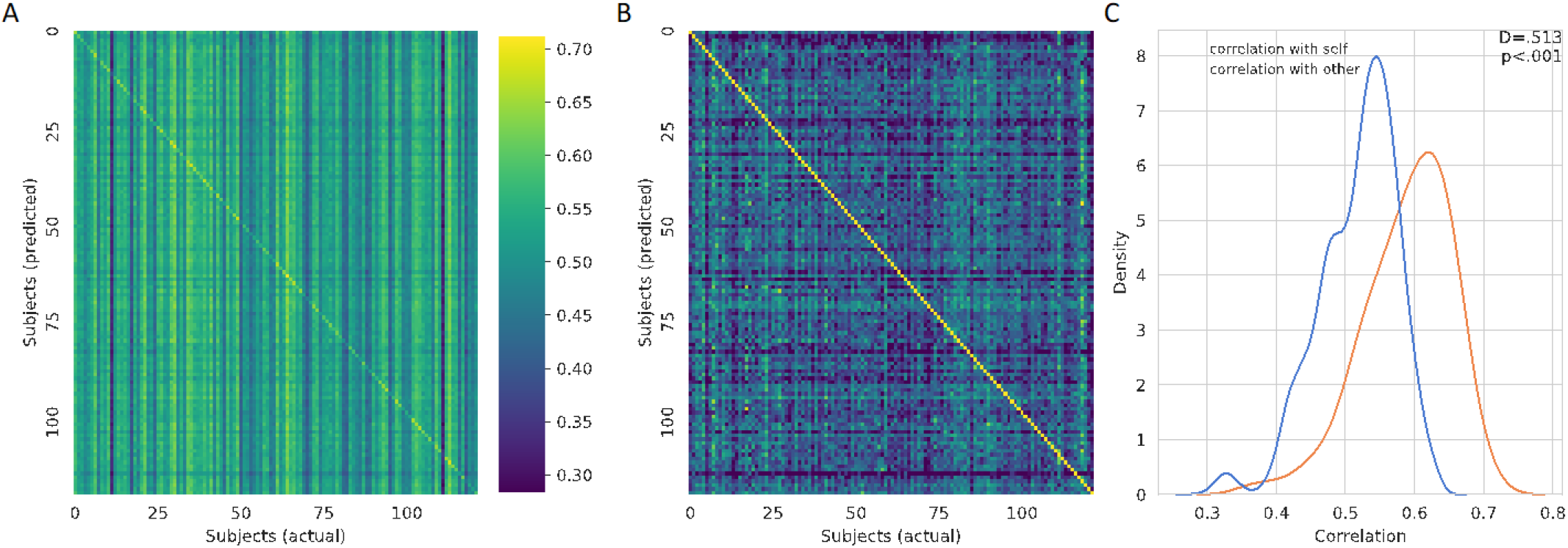
(**A**) Overall correlation matrix for all tasks. The predicted maps (y-axis) were compared to the actual maps (x-axis) for all of the subjects. The visible diagonal indicates that the predicted maps were more correlated with their own actual maps than the maps of other subjects. (**B**) Row and column normalized correlation matrix to remove mean correlation. (**C**) Distribution of the diagonal elements of the (un-normalized) correlation matrix in orange and the extra-diagonal elements in blue visualized using a kernel density estimation and overlapping normalized histogram. A Kolmogorov-Smirnov test between the two distributions gives a highly significant difference, *p* < .001, indicating that the predicted task maps are more correlated to their actual maps than the maps of other subjects.

**Figure 4.**
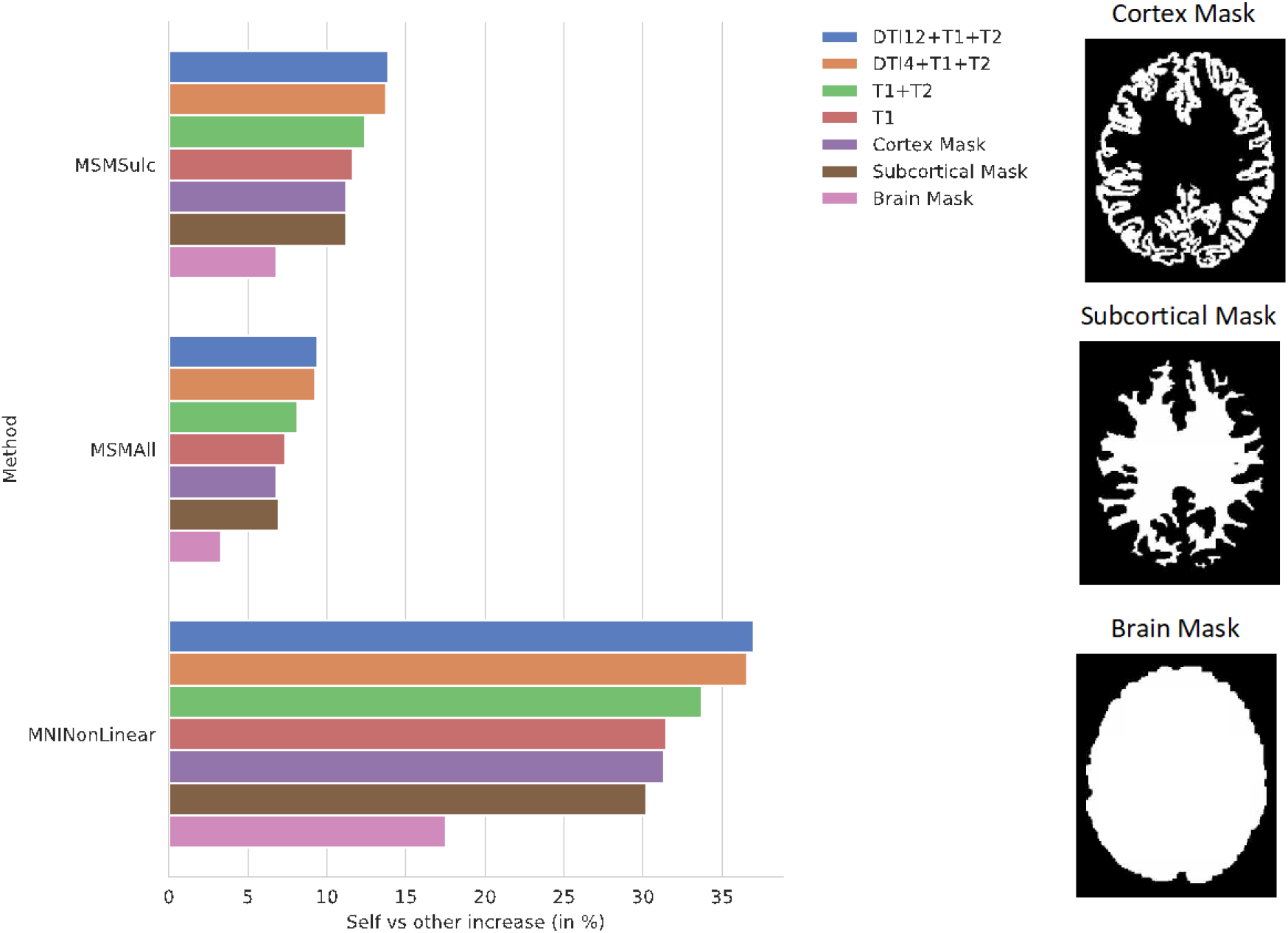
Ablation Study. Self (matrix diagonal) vs other (extra-diagonal elements) correlation in predictions shows the difference between average correlation of the predictions to the actual maps and the average correlation of the predictions to the actual maps of all the other subjects. Ablation study results are shown for the CNN models using the proposed combination of the anatomical imaging and DTI data for each of the 3 b-values (T1+T2+DTI12), the anatomical imaging and overall DTI data (T1+T2+DTI4), the anatomical imaging alone (T1+T2), and the T1w imaging alone (T1) as well as masked images of the cortex (Cortex Mask), subcortical areas (Subcortical Mask), and the whole brain (Brain Mask). Examples of the masked images for a single subject are shown on the right. Results were compared using the MSMSulc and the MSMAll registered template surfaces as well as non-linear registration into MNI volume space. Compared to the MSMSulc registered template surfaces, the MSMAll registered templates did account for some of the predicted variation, but the predicted maps were still more correlated with their own actual maps than all other actual maps on average for all of the input types and methods. Comparing the predictions to the actual maps non-linearly warped into MNI volume space resulted in a much higher increase correlation between the predictions and actual maps than comparing the maps using the surface templates. The models trained on the masked images were able to predict individual subject variation, but the model using the masked brain images exhibited a marked decrease in performance.

The model was also able to predict the lateralization of task function. The predicted lateralization indices were significantly correlated (*p* < .001) with the actual lateralization indices for 28 out of the 47 activation maps. The linear regression of the lateralization indices between the predicted and actual maps for the exemplary tasks is shown in *Figure 5*.

**Figure 5.**
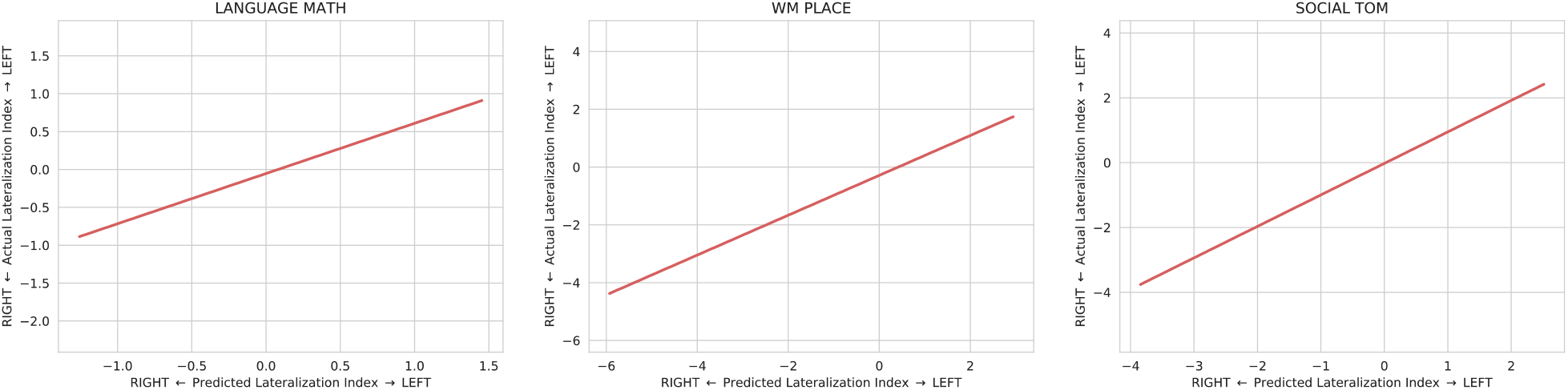
Linear regression fit with 95% confidence intervals between the predicted and actual lateralization indices for the LANGUAGE MATH, WM PLACE, and SOCIAL TOM activation maps. The lateralization index is defined as the difference between the left hemisphere and right hemisphere at the peak location for the predicted map. All three plots show that the predicted lateralization indices are positively correlated with the with the actual lateralization indices indicating that the predicted maps are able to capture some of the variation of lateralization seen in the actual maps.

The ablation study shows the correlation of the model using all of the proposed structural data plotted against models trained using DTI data not computed at separate b-values, T1w with T2w data, and T1w data only as well as models trained on binary mask images of either the cortex, subcortical matter, or the whole brain (*Figure 4*). All models were able to predict individual subject variation, but the model using all of the data performed the best. This indicates that the information contained in the additional imaging features and features improved the model’s prediction. The model trained on the mask of the brain resulted in the least increase in correlation over the average.

The test-retest reliability of the predicted maps as measured by ICC was greater for all domains than the actual maps (*Figure S7*) and the ICC of all predicted maps on average was .95 compared to .61 for the actual maps.

## Discussion

We demonstrated that structural imaging contains information that is predictive of inter-subject variations in task activations using a CNN. By contrast, Tavor et al. did not find structural imaging features to be predictive of variations in task activations when using a simple linear regression model (Tavor et al., 2016). CNNs, however, are able to extract rich and complex features that give them an immense advantage, with enough data, over linear regression. Therefore, the CNN was able to extract features from the imaging that predicted task activations over a wide array of domains that correlated well to the actual tfMRI activations. Previous research has not reported such a broad correlation between structural imaging and individual-level variations in functional activation, although some research has reported the connectivity of reconstructed white matter tracts to be predictive of visual tasks (Saygin et al., 2016). In contrast, our model was able to predict variations in task activation in 47 different activation maps across seven different task domains (motor, language, working memory, relational, emotion, social, and gambling). We demonstrated that our model was able to predict deviations from the group average activation (Figure 2) as well as variations in lateralization (Figure 5). Even when alignment between subjects was performed using MSMAll that takes into account functional mapping features, our model was still able to predict variation in the task activation of individual subjects (*Figure S6*). Therefore, it is not likely that the variation captured by the predictions is solely the result of the functional misalignment between subjects.

The ablation study showed that adding DTI along with T1w and T2w anatomical imaging resulted in better predictions than the T1w and T2w model alone (*Figure 5*). Thus, the microstructural diffusion imaging features contain information not encompassed by the anatomical imaging that is predictive of individual-level variation in task activations. Surprisingly, the ablation study also showed that the shape of either the sub-cortex alone or the cortex alone was sufficient to predict individual-level variation using a CNN. However, when the model was only able to see the shape of the brain mask, the model was not as accurate. The information present in the shape of the sub-cortex but not in the shape of the brain is the structure and location of the gyri and sulci, also known as the cortical folding patterns. The model was able to use the shape of the sub-cortex to infer the cortical folding patterns and then use this information to predict the task activation patterns throughout the cortex. Without these patterns, the model is unable to predict variations between individual subjects to the same extent as delineated by the poorer performance of the model using only the brain mask. This finding along with previous research showing that cortical folding patterns are unique to the individual (Duan et al., 2020), influenced by the tension from brain connections (David C Van Essen, 2020), and correlated to behavior (Whittle et al., 2009) as well as neuropsychological impairments (Shaw et al., 2012) indicates that folding patterns may be an integral part to how brain function is derived from brain structure.

Structural imaging is immune to some of the sources of noise that make tfMRI mapping unreliable, such as neurovascular uncoupling, poor task performance, and cardiac rhythm. Therefore, it is not surprising that the predictions were more reliable than the actual tfMRI maps when evaluated in subjects that were scanned twice (*Figure S7*). However, it should be noted that more thorough processing of the fMRI to remove structured noise components could also improve the reliability of the fMRI signal (Glasser et al., 2018; Parkes, Fulcher, Yücel, & Fornito, 2018). More dependable functional activation maps would greatly assist clinicians in cases where task activation mapping is critical for patient care, including neurosurgical operative planning. Accurate task mapping predictions could give neurosurgeons the ability to visualize eloquent areas and avoid potential surgically induced deficits, even without collecting any fMRI data. This would be particularly advantageous in situations where collecting fMRI data is infeasible due to limitations in patient performance or insufficient resources.

## Conclusion

Structural imaging, when paired with a CNN, was predictive of inter-subject variations in 47 different task activation maps across seven task domains. These findings suggest that anatomical and microstructural features contain information that is predictive of unique functional brain activations in individuals.

## Acknowledgments

This work was completed utilizing the Holland Computing Center of the University of Nebraska, which receives support from the Nebraska Research Initiative. Data were provided by the Human Connectome Project, WU-Minn Consortium (Principal Investigators: David Van Essen and Kamil Ugurbil; 1U54MH091657), funded by the 16 NIH Institutes and Centers that support the NIH Blueprint for Neuroscience Research; and by the McDonnell Center for Systems Neuroscience at Washington University.

## Author Contributions

D.E. and M.A. contributed to study conception, design, and drafting of the manuscript. D.E. performed the analysis and interpretation of the data.

## Compliance with Ethical Standards

All procedures performed in studies involving human participants were in accordance with the ethical standards of the institutional and/or national research committee and with the 1964 Helsinki declaration and its later amendments or comparable ethical standards.

## References

Barch, D. M., Burgess, G. C., Harms, M. P., Petersen, S. E., Schlaggar, B. L., Corbetta, M.,… Feldt, C. (2013). Function in the human connectome: task-fMRI and individual differences in behavior. Neuroimage, 80, 169–189.

Coalson, T. S., Van Essen, D. C., & Glasser, M. F. (2018). The impact of traditional neuroimaging methods on the spatial localization of cortical areas. Proceedings of the National Academy of Sciences, 115(27), E6356–E6365. doi:10.1073/pnas.1801582115

Duan, D., Xia, S., Rekik, I., Wu, Z., Wang, L., Lin, W.,… Li, G. (2020). Individual identification and individual variability analysis based on cortical folding features in developing infant singletons and twins. Human Brain Mapping, 41(8), 1985–2003.

Elliott, M. L., Knodt, A. R., Ireland, D., Morris, M. L., Poulton, R., Ramrakha, S.,… Hariri, A. R. (2020). What Is the Test-Retest Reliability of Common Task-Functional MRI Measures? New Empirical Evidence and a Meta-Analysis. Psychological Science, 0(0), 0956797620916786. doi:10.1177/0956797620916786

Ellis, D. G., White, M. L., Hayasaka, S., Warren, D. E., Wilson, T. W., & Aizenberg, M. R. (2020). Accuracy analysis of fMRI and MEG activations determined by intraoperative mapping. Neurosurgical Focus, 48(2), E13.

Ganepola, T., Nagy, Z., Ghosh, A., Papadopoulo, T., Alexander, D. C., & Sereno, M. I. (2018). Using diffusion MRI to discriminate areas of cortical grey matter. NeuroImage, 182, 456–468. doi:https://doi.org/10.1016/j.neuroimage.2017.12.046

Garyfallidis, E., Brett, M., Amirbekian, B., Rokem, A., Van Der Walt, S., Descoteaux, M., & Nimmo-Smith, I. (2014). Dipy, a library for the analysis of diffusion MRI data. Frontiers in Neuroinformatics, 8(8). doi:10.3389/fninf.2014.00008

Glasser, M. F., Coalson, T. S., Bijsterbosch, J. D., Harrison, S. J., Harms, M. P., Anticevic, A.,… Smith, S. M. (2018). Using temporal ICA to selectively remove global noise while preserving global signal in functional MRI data. Neuroimage, 181, 692–717.

Glasser, M. F., Coalson, T. S., Robinson, E. C., Hacker, C. D., Harwell, J., Yacoub, E.,… Jenkinson, M. (2016). A multi-modal parcellation of human cerebral cortex. Nature, 536(7615), 171–178. Retrieved from https://www.ncbi.nlm.nih.gov/pmc/articles/PMC4990127/pdf/emss-68870.pdf

Glasser, M. F., Sotiropoulos, S. N., Wilson, J. A., Coalson, T. S., Fischl, B., Andersson, J. L.,… Consortium, W. U.-M. H. (2013). The minimal preprocessing pipelines for the Human Connectome Project. Neuroimage, 80, 105–124. doi:10.1016/j.neuroimage.2013.04.127

He, K., Zhang, X., Ren, S., & Sun, J. (2016). Identity mappings in deep residual networks. Paper presented at the European conference on computer vision.

Myronenko, A. (2018). 3D MRI brain tumor segmentation using autoencoder regularization. Paper presented at the International MICCAI Brainlesion Workshop.

Osher, D. E., Saxe, R. R., Koldewyn, K., Gabrieli, J. D. E., Kanwisher, N., & Saygin, Z. M. (2015). Structural Connectivity Fingerprints Predict Cortical Selectivity for Multiple Visual Categories across Cortex. Cerebral Cortex, 26(4), 1668–1683. doi: 10.1093/cercor/bhu303

Parkes, L., Fulcher, B., Yücel, M., & Fornito, A. (2018). An evaluation of the efficacy, reliability, and sensitivity of motion correction strategies for resting-state functional MRI. Neuroimage, 171, 415–436.

Ronneberger, O., Fischer, P., & Brox, T. (2015). U-net: Convolutional networks for biomedical image segmentation. Paper presented at the International Conference on Medical image computing and computer-assisted intervention.

Saygin, Z. M., Osher, D. E., Koldewyn, K., Reynolds, G., Gabrieli, J. D., & Saxe, R. R. (2012). Anatomical connectivity patterns predict face selectivity in the fusiform gyrus. Nature Neuroscience, 15(2), 321. Retrieved from https://www-nature-com.library1.unmc.edu/articles/nn.3001.pdf

Saygin, Z. M., Osher, D. E., Norton, E. S., Youssoufian, D. A., Beach, S. D., Feather, J.,… Kanwisher, N. (2016). Connectivity precedes function in the development of the visual word form area. Nature Neuroscience, 19(9), 1250–1255. Retrieved from https://www.ncbi.nlm.nih.gov/pmc/articles/PMC5003691/pdf/nihms801288.pdf

Shaw, P., Malek, M., Watson, B., Sharp, W., Evans, A., & Greenstein, D. (2012). Development of cortical surface area and gyrification in attention-deficit/hyperactivity disorder. Biological Psychiatry, 72(3), 191–197.

Shrout, P. E., & Fleiss, J. L. (1979). Intraclass correlations: uses in assessing rater reliability. Psychological Bulletin, 86 2, 420–428.

Sotiropoulos, S. N., Jbabdi, S., Xu, J., Andersson, J. L., Moeller, S., Auerbach, E. J.,… Jenkinson, M. (2013). Advances in diffusion MRI acquisition and processing in the Human Connectome Project. Neuroimage, 80, 125–143. Retrieved from https://www.ncbi.nlm.nih.gov/pmc/articles/PMC3720790/pdf/nihms483085.pdf

Tavor, I., Jones, O. P., Mars, R., Smith, S., Behrens, T., & Jbabdi, S. (2016). Task-free MRI predicts individual differences in brain activity during task performance. Science, 352(6282), 216–220.

Van Essen, D. C. (2020). A 2020 view of tension-based cortical morphogenesis. Proceedings of the National Academy of Sciences, 117(52), 32868–32879.

Van Essen, D. C., Ugurbil, K., Auerbach, E., Barch, D., Behrens, T. E., Bucholz, R.,… Yacoub, E. (2012). The Human Connectome Project: a data acquisition perspective. Neuroimage, 62(4), 2222–2231. doi:10.1016/j.neuroimage.2012.02.018

Weng, H.-H., Noll, K. R., Johnson, J. M., Prabhu, S. S., Tsai, Y.-H., Chang, S.-W.,… Liu, H.-L. (2018). Accuracy of Presurgical Functional MR Imaging for Language Mapping of Brain Tumors: A Systematic Review and Meta-Analysis. Radiology, 286(2), 512–523. doi: 10.1148/radiol.2017162971

Whittle, S., Allen, N. B., Fornito, A., Lubman, D. I., Simmons, J. G., Pantelis, C., & Yücel, M. (2009). Variations in cortical folding patterns are related to individual differences in temperament. Psychiatry Research: Neuroimaging, 172(1), 68–74.

Woolrich, M. W., Ripley, B. D., Brady, M., & Smith, S. M. (2001). Temporal autocorrelation in univariate linear modeling of FMRI data. Neuroimage, 14(6), 1370–1386. doi:10.1006/nimg.2001.0931

Wu, Y., & He, K. (2018). Group normalization. Paper presented at the Proceedings of the European Conference on Computer Vision (ECCV).

