## Supplement Figures and Table for "Structural brain imaging predicts individual-level task activation maps using deep learning"

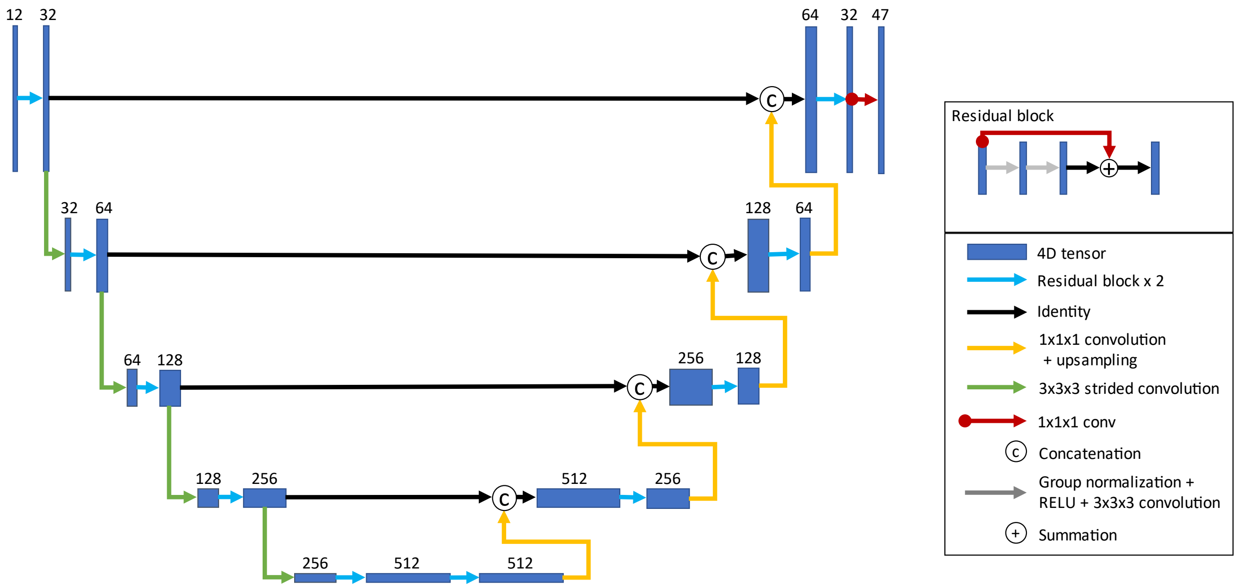


*Figure S1* - **Model architecture.** A U-Net architecture was used with two residual blocks for each encoding and decoding layer. The inputs for each consecutive encoding layer are downsampled using a strided convolution. The decoding layer takes as inputs the outputs from the last encoding layer. Each consecutive decoding layer concatenates the outputs from the encoding layer at the same resolution as well as upsampled outputs from the previous decoding layer. A 1x1x1 convolution linearly resamples the 32 channel outputs from final decoding layer into 47 volumes, one for each task activation map.

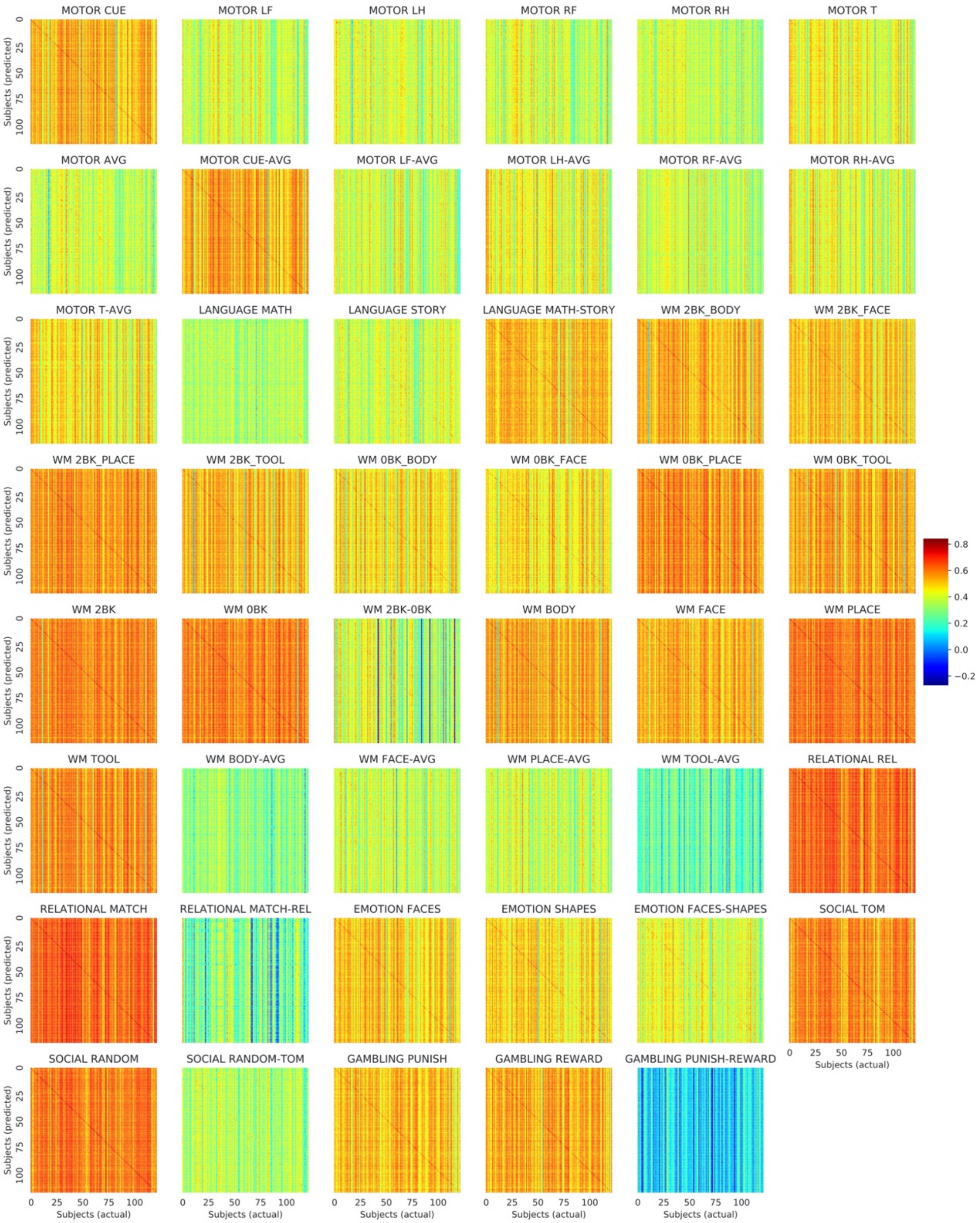


*Figure S2* - Correlation matrices between the predicted task activation maps on the y-axis and the actual activation maps on the x-axis. A visible diagonal indicates that the predicted activation maps are more correlated with the respective actual activation map than with the activation maps of other subjects.

CUE=task cue; LF=left foot; LH=left hand; RF=right foot; RH=right hand; T=tongue; AVG=average; WM=Working Memory; 0BK=0 Back; 2BK=2 Back.

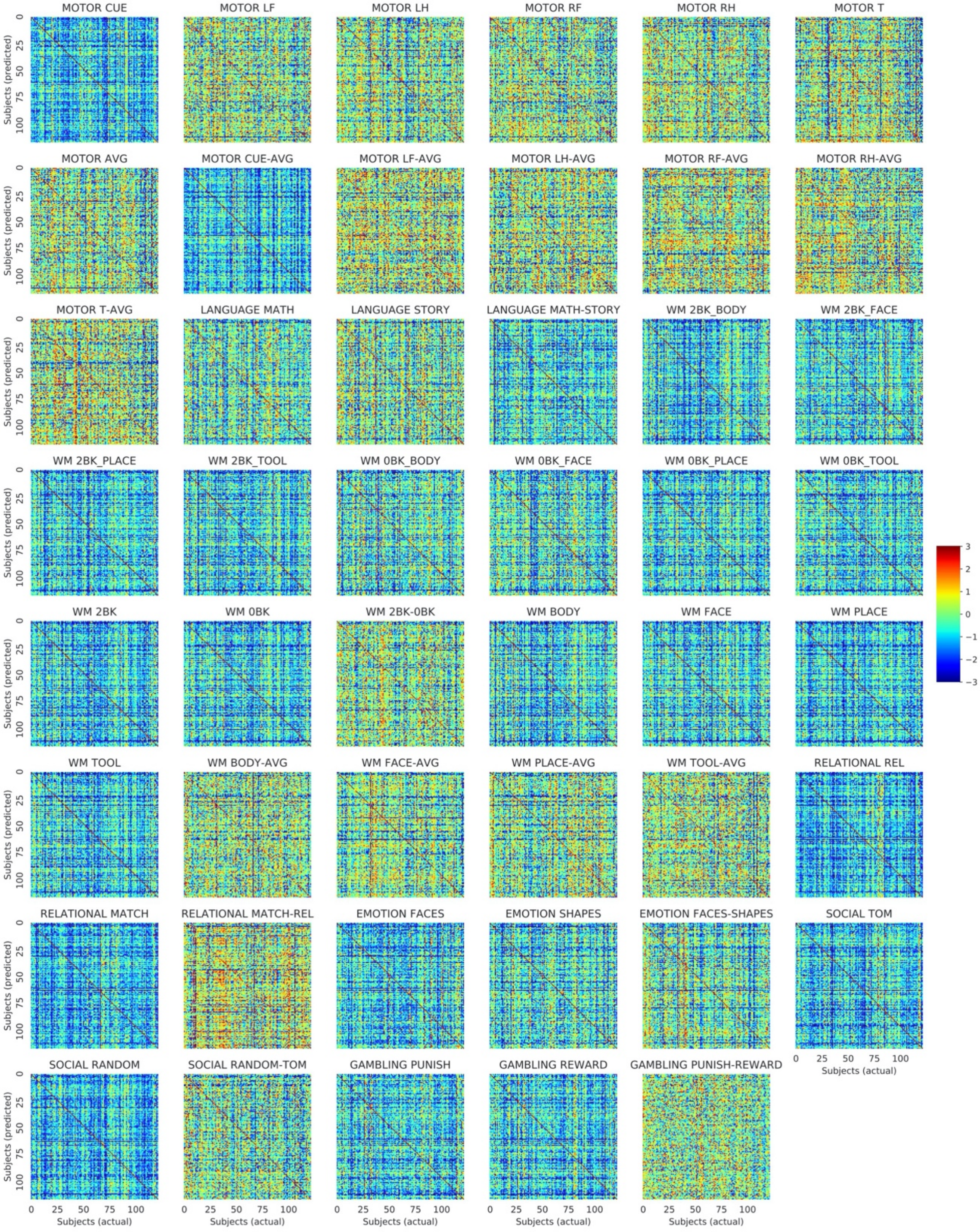


*Figure S3* - Correlation matrices of the individual task maps that have been row and column normalized to account for the increased variability in the actual activation maps than in the predicted activation maps. The diagonal indicating the correlation between the predicted and actual maps is even more apparent after normalization.

CUE=task cue; LF=left foot; LH=left hand; RF=right foot; RH=right hand; T=tongue; AVG=average; WM=Working Memory; 0BK=0 Back; 2BK=2 Back.

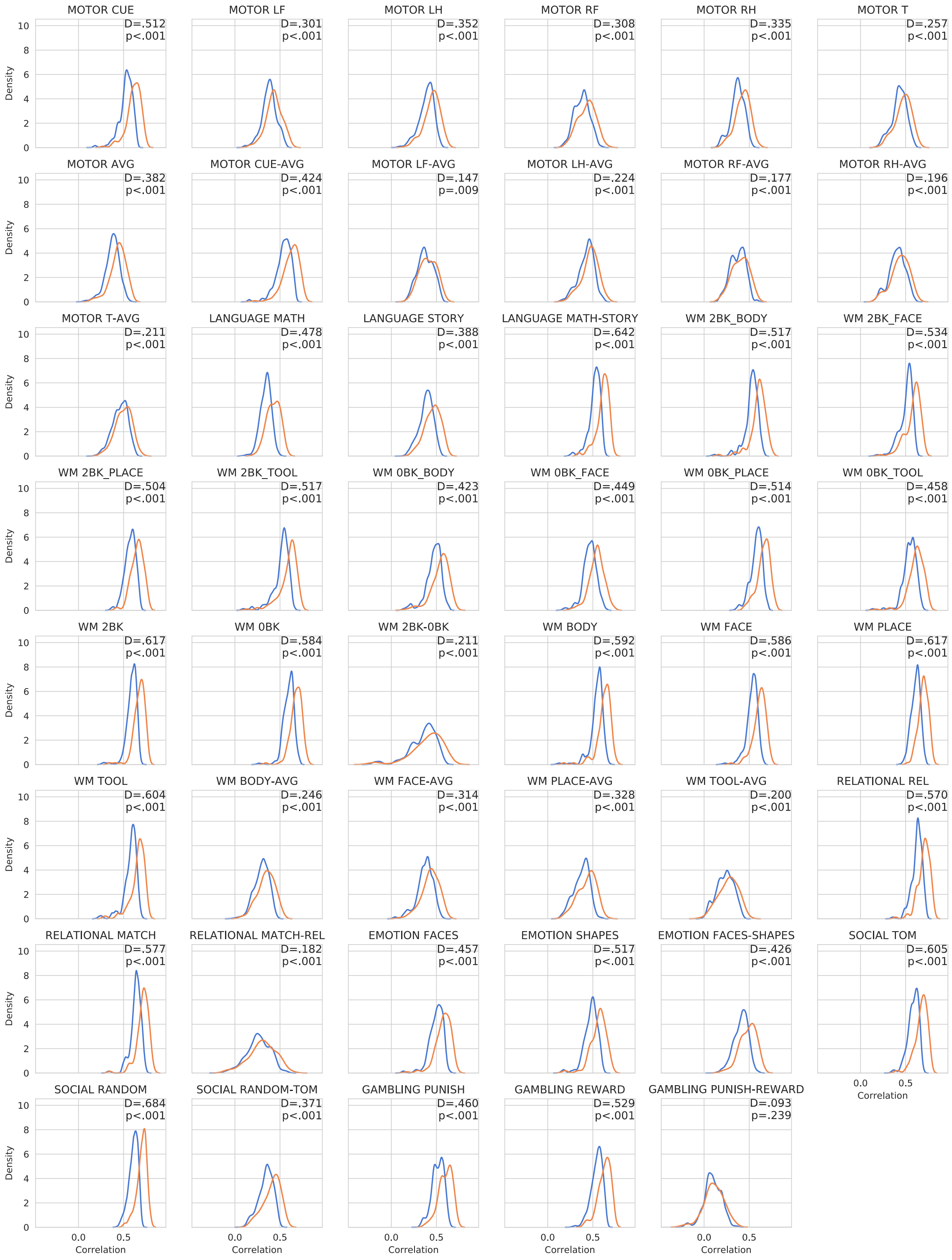


*Figure 4* - Histogram comparing the correlation of the predicted individual motor and language task maps for a subject to the actual map for that subject in orange and the correlation between the predicted maps and the actual task activation maps for the other subjects in blue.

CUE=task cue; LF=left foot; LH=left hand; RF=right foot; RH=right hand; T=tongue; AVG=average; WM=Working Memory; 0BK=0 Back; 2BK=2 Back.

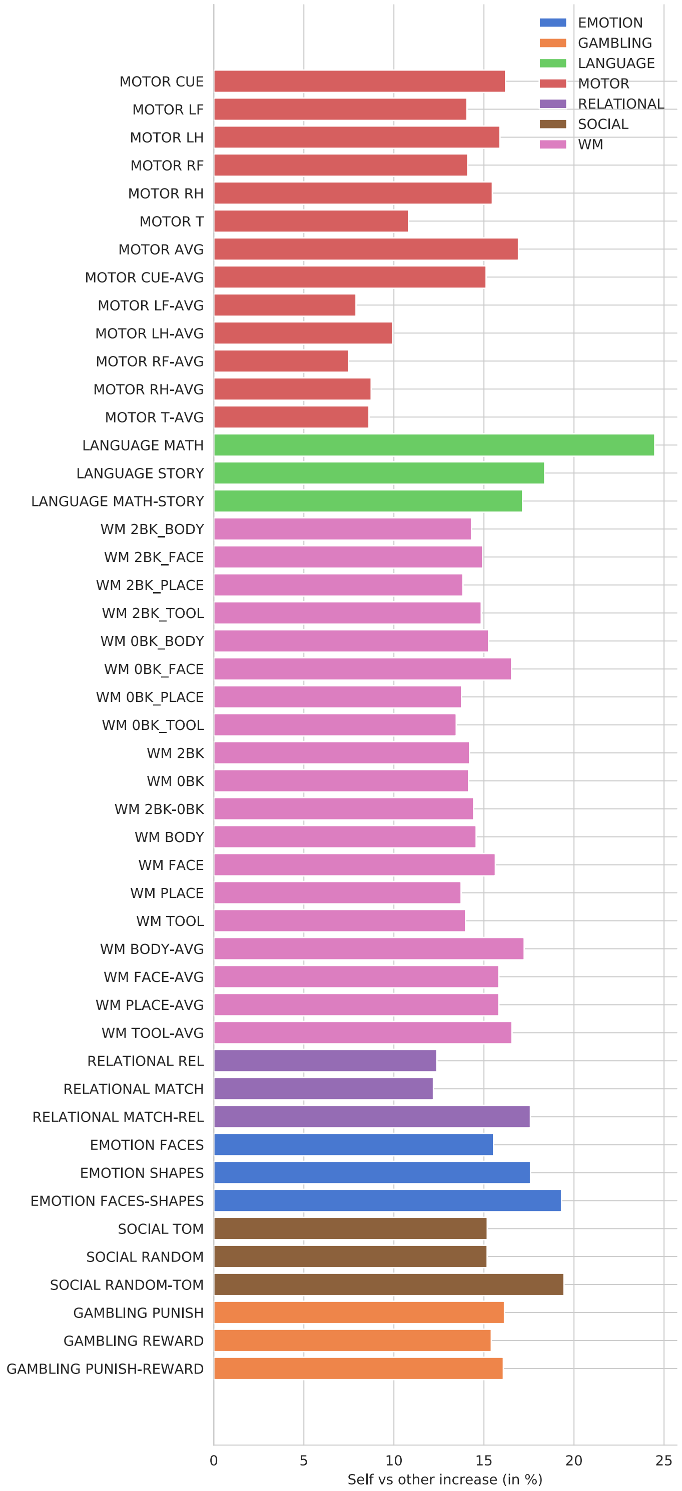


*Figure S5* - **Self vs Other increase**. The difference between the average correlation between the predicted and actual maps (diagonal elements) and the average correlation between the predicted maps and the maps of all the other subjects (extra-diagonal elements) as a percentage relative to the average of the extra- diagonal elements. The positive values show that on average, the predictions match the actual maps (self) better than the average of the extra-diagonal elements (others).

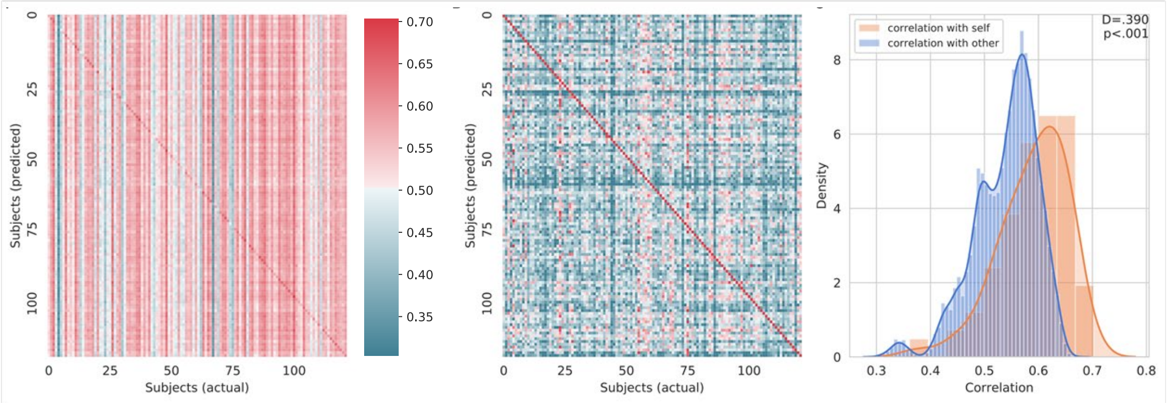


*Figure S6* - (**A**) Average correlation matrix for all tasks compared using MSMAll registered surface templates rather than the MSMSulc registered surface templates. The predicted maps (y-axis) were compared to the actual maps (x-axis) for all of the subjects. The MSMAll algorithm attempts to minimize individual subject variations in functional localization by using resting state fMRI and structural features for surface registration. Even with the MSMAll registration, the visible diagonal indicates that the predicted maps were more correlated with their own actual maps than the maps of other subjects. (**B**) Row and column normalized correlation matrix to remove mean correlation. The normalized correlation matrix is heavily diagonal dominant. (**C**) Distribution of the diagonal elements of the (un-normalized) correlation matrix in orange and the extra-diagonal elements in blue visualized using a kernel density estimation and overlapping normalized histogram. A Kolmogorov-Smirnov test between the two distributions gives a highly significant difference $p < .001$. This indicates individual variances in the task activation maps that the model is able to accurately predict are not solely the result of minor functional misalignments between subjects, as much of the variation was still predicted by the model even after the MSMAll surface templates corrected for the functional misalignment via registration.

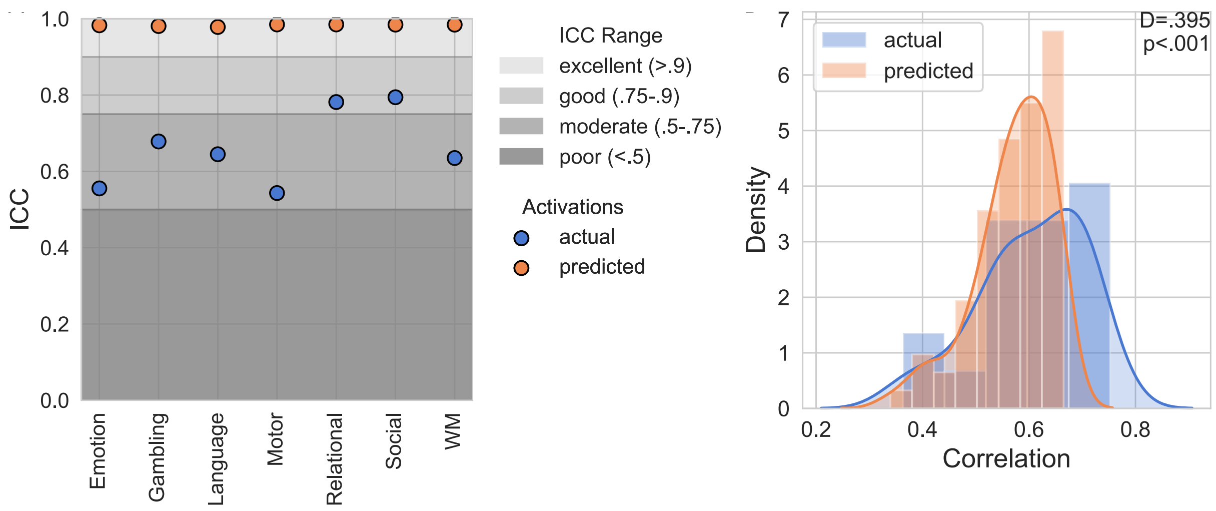


*Figure S7* - **Test-retest analysis**. (**A**) Reliability of the predicted activation maps compared to that of the actual task-fMRI activation maps. Reliability was determined by assessing the intraclass correlation (ICC) of the maps over the entire cortex for subjects who were scanned and then rescanned four months later. The predicted maps had excellent ICC scores for all domains that were higher than the ICC scores of the actual task-fMRI maps. (**B**) Histogram and kernel density estimation for the test-retest correlation of actual task-fMRI maps to themselves in blue compared to the correlation of the predicted maps to the actual task-fMRI maps from the opposite test-retest session in orange. The predicted maps were not as correlated to the test-retest maps as the actual maps. Together these results show that the predicted maps were highly consistent but were not able to predict all the information that is captured by task fMRI.

| Domain | Attributes | Activation Maps |
|---|---|---|
| Emotion | Shape matching compared to face matching with angry or fearful expressions | FACES, SHAPES, FACES-SHAPES |
| Gambling | Incentive processing, punishments, and rewards | PUNISH, REWARD, PUNISH-REWARD |
| Language | Auditory and phonological stories and arithmetic | STORY, MATH, MATH-STORY |
| Motor | Hand, foot, and tongue movements | CUE, LF, LH, RF, RH, T, AVG, CUE-AVG, LF-AVG, LH-AVG, RF-AVG, RH-AVG, T-AVG |
| Relational | Matching shapes and textures | REL, MATCH, MATCH-REL |
| Social | Random interactions compared to social interactions | TOM, RANDOM, RANDOM-TOM |
| Working Memory | N-back working memory, faces, places, tools, and body parts | 2BK BODY, 2BK FACE, 2BK PLACE, 2BK TOOL, 0BK BODY, -0BK FACE, 0BK PLACE, 0BK TOOL, 2BK-0BK, BODY, FACE, PLACE, TOOL, BODY-AVG, FACE-AVG, PLACE-AVG, TOOL-AVG |

*Table S1* – Nomenclature and attributes for the task domains and activation maps included in this paper. The HCP uses the same nomenclature in their data releases and publications.
